## Supplementary data for "Bacterial filamentation is an in vivo mechanism for cell-to-cell spreading"

### SUPPLEMENTARY ITEMS

#### Supplementary Text

Taxonomic summary of *Bordetella atropi*

***Bordetella atropi* defines a new species.** This bacterium was originally found in its nematode host, *Oscheius tipulae* strain JU1501*.* It is orally transmitted, with no evidence of vertical transmission. The bacterium was cultured in pure culture in vitro in Luria Broth (LB) and other standard bacterial media and was amenable to conjugation with *E. coli* for transmission of plasmids. Phylogenomic analysis based on 92 highly conserved bacterial gene sequences placed this bacterium in the *Bordetella* genus as a new species.

**Life cycle and symptoms in its host.** Symptoms of infection were originally discovered by Nomarski light microscopy as small coccobacilli in the intestinal lumen and inside intestinal cells. The infection was passed from uninfected animals to infected animals through co-culture on the same plate, suggesting a fecal-oral route of transmission. Additionally, since the bacterium was isolated and cultured in vitro, animals were reinfected orally through exposure to the bacterium on a plate for as little as 2 hours. Growth in LB resulted in mostly coccobacilli bacteria, with a small proportion of the population displaying flagellar motility. It is likely that the coccobacillus form of the bacterium is infectious to *O. tipulae.*

Observations of infection from fluorescent in situ hybridization (FISH) experiments using specific 16S rRNA probes and transmission electron microscopy (TEM) indicated that *B. atropi* has a filamentous morphology during early infection, which preceded the observation of hundreds to thousands of coccobacilli in late infection. The filamentous form was often observed to be simultaneously infecting intestinal cells and indicating invasion of neighboring cells. All infection phenotypes were only observed in intestinal cells, except for coccobacilli in the intestinal lumen. All post-embryonic stages showed signs of infection, including non-feeding dauer animals (if exposed prior to entering dauer).

The type strain (LUAb4) was isolated from a rotting apple below a wild apple tree near Kerarmel, Plouezoc'h, France (GPS coordinates 48.65151, -3.85573) on August 6, 2008.

The etymology of the type species name *B. atropi* is based on the filamentation phenotype (long threads) and the high level of pathogenicity to the cognate *O. tipulae* strain JU1501. This species was named after Atropos, the Greek Fate who cuts the thread of life.

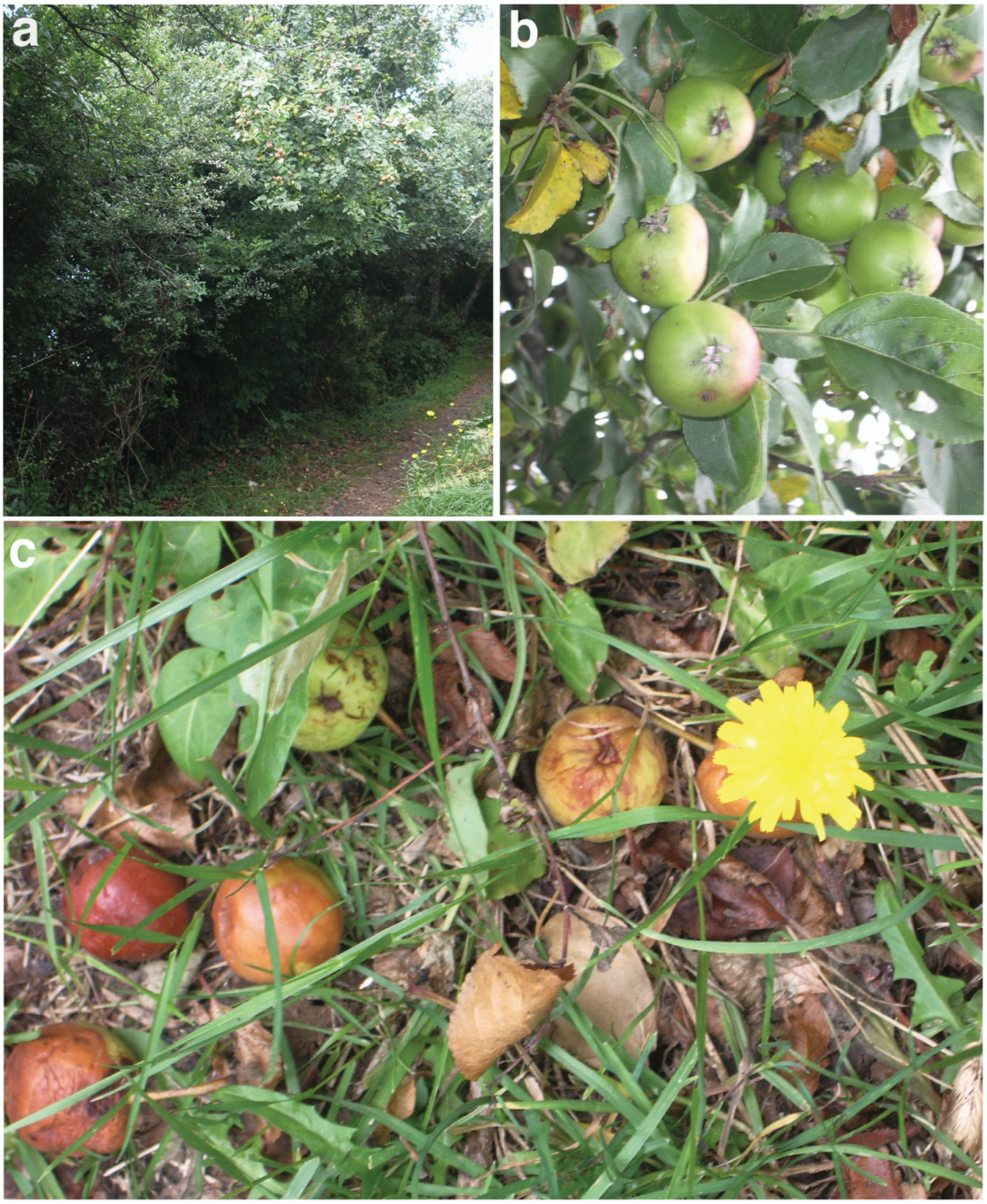


Supplementary Fig. 1**. Rotting European crab apples sampled in Kerarmel, Plouezoc'h, France. a** A wild crab apple tree with **b** ripening fruit on the stem and **c** rotting fruit below. The rotting apples were 23-25 mm in width and were taken for wild nematode sampling. *O. tipulae* strain JU1501 was isolated from these samples.

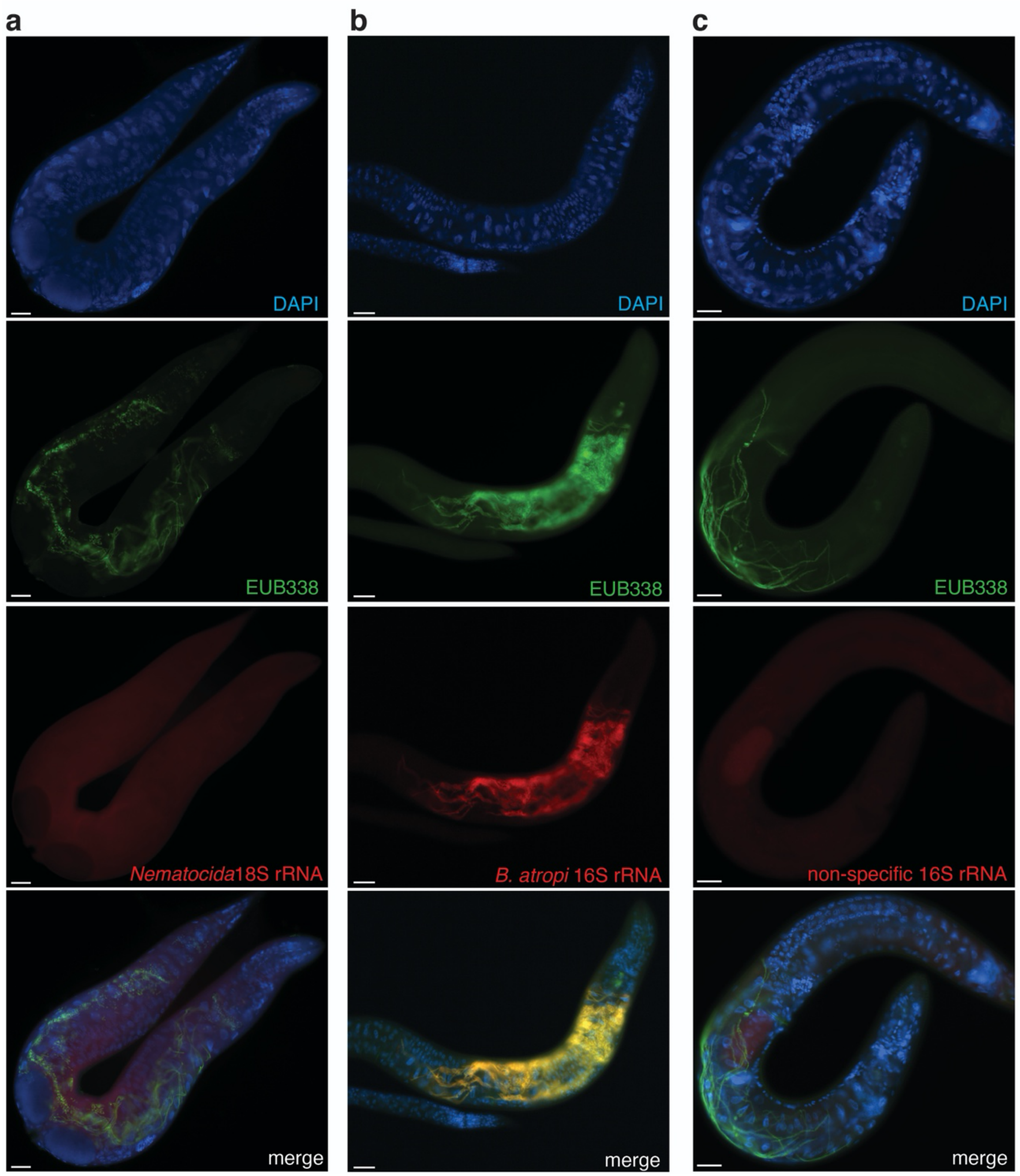


Supplementary Fig. 2. **Fluorescent micrographs of wild *O. tipulae* strain JU1501 infected with *B. atropi*.** Animals were stained by DAPI, a universal bacterial 16S rRNA probe labeled with FAM (EU338), and FISH probes labeled with CF610 specific to the small subunit of microbial rRNA, either **a** *Nematocida* 18S probes microA, microC, microE, **b** *B. atropi* 16S probe b004, or **c** Alphaproteobacteria 16S probe b002. Scale bars are 25 μm.

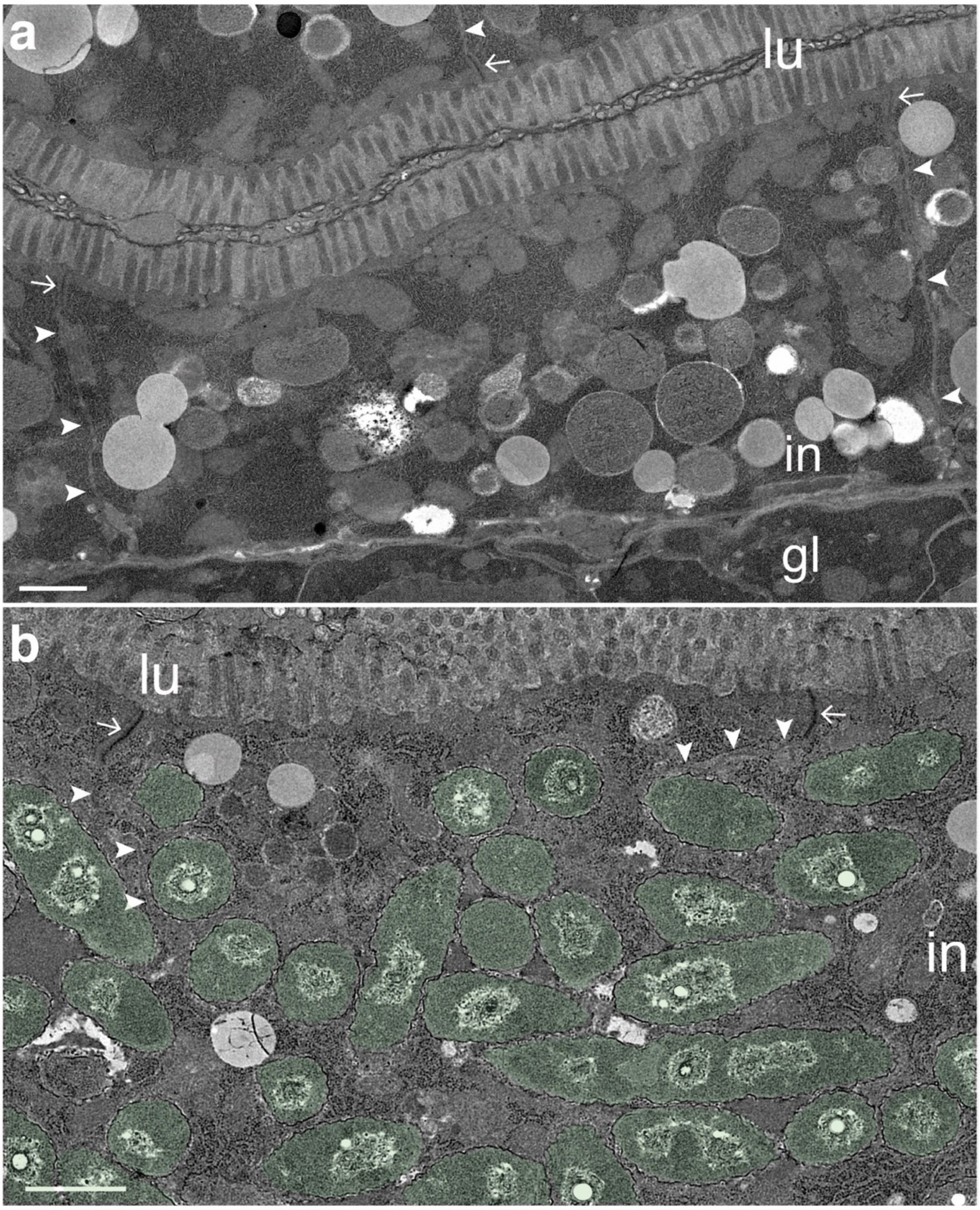


Supplementary Fig. 3. **TEM images of *B. atropi* phenotypes in *O. tipulae*. a** Uninfected and **b** infected intestine at 48 hpi showing the lumen (*lu*) with electron dense apical junctions (*arrows*) followed by the lateral intestinal membranes (*arrowheads*). The germline is indicated (*gl*). Scalebars are 1 μm.

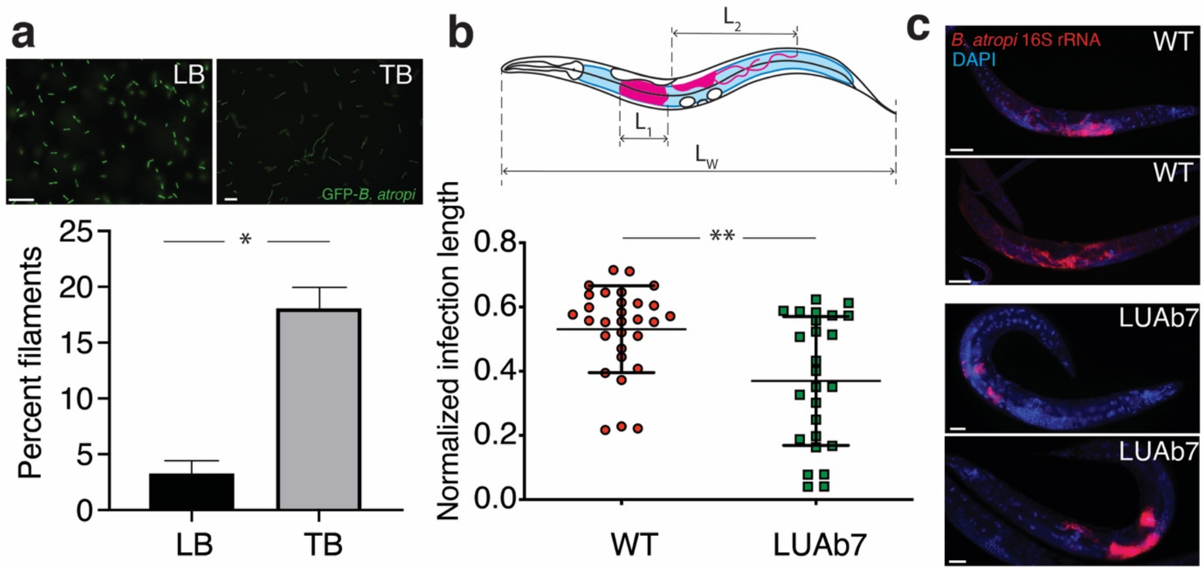


Supplementary Fig. 4. **Filamentation of *B. atropi* in TB and anterior-posterior (A-P) spreading in vivo. a** GFP-*B. atropi* was grown in LB ON and was transferred to LB or TB at 32°C for 48 hours. Bacteria were binned as filaments if greater than 4 mm in length, representing >4 bacterial cell lengths. The mean and SD of two independent trials is shown with p=0.011 (*) by the unpaired two-tail t test. Representative images are shown on top. Scale bars are 10 μm. **b** Animals were infected for 34 hours and stained with FISH. A schematic for describing A-P spreading (top), where the length of each contiguous infection in an animal was measures along the A-P axis ($L_1$, $L_2$,…,$L_N$) and summed. This total infection length was normalized to the A-P length of the animal ($L_W$) giving the normalized infection length (*top*). Results are combined from 2 independent replicates, means and SD are shown, p=0.002 (**) by the Mann Whitney two-tail t test (*bottom*). **c** Representative images from **b** are shown. Scale bars are 20 μm.

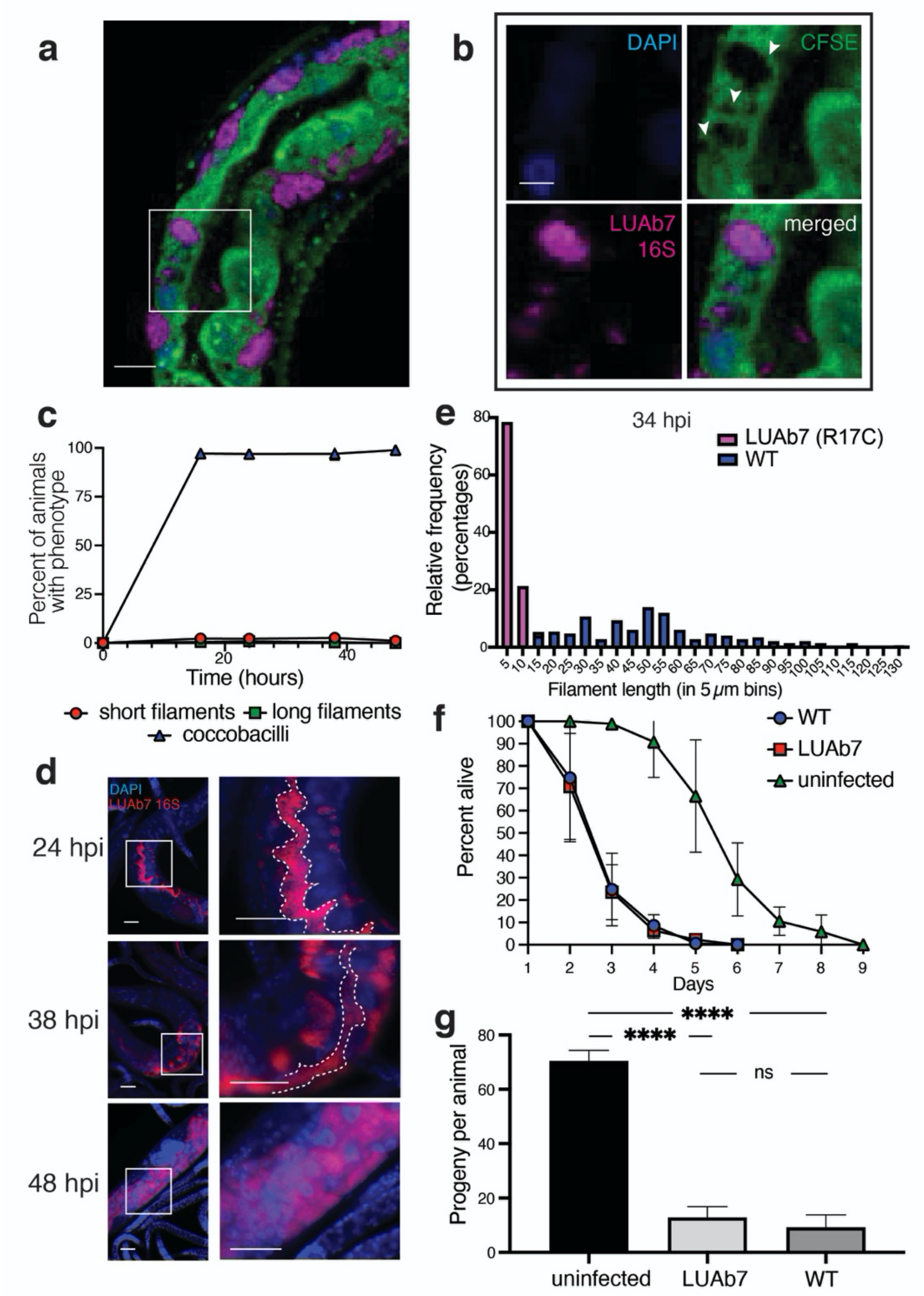


Supplementary Fig. 5. **Characterization of LUAb7 in vivo phenotypes. a** Representative confocal image of an animal infected with LUAb7. **b** Inset of region indicated by white box in **a** showing the overlap of CFSE-fluorescence clearing (arrowheads) with FISH signal from the *B. atropi*-specific 16S probe. **c** Pulse chase infection time course of two independent replicates is shown for LUAb7, n > 200 animals for each time point in 2 independent replicates. **d** Representative images of phenotypes at indicated time points in a. Dashed lines delineate the lumen. **e** Distribution of in vivo filament lengths of LUAb7 compared to WT, n = 30 animals in 2 independent replicates. **f** Life span of animals infected with either WT or LUAb7 compared to uninfected animals, n = 40 animals in 2 independent replicates. **g** Broodsize of animals infected with LUAb7 compared to WT and uninfected animals, n = 40 animals in 2 independent replicates. Scale bars are 5 μm in **b** and 10 μm elsewhere.

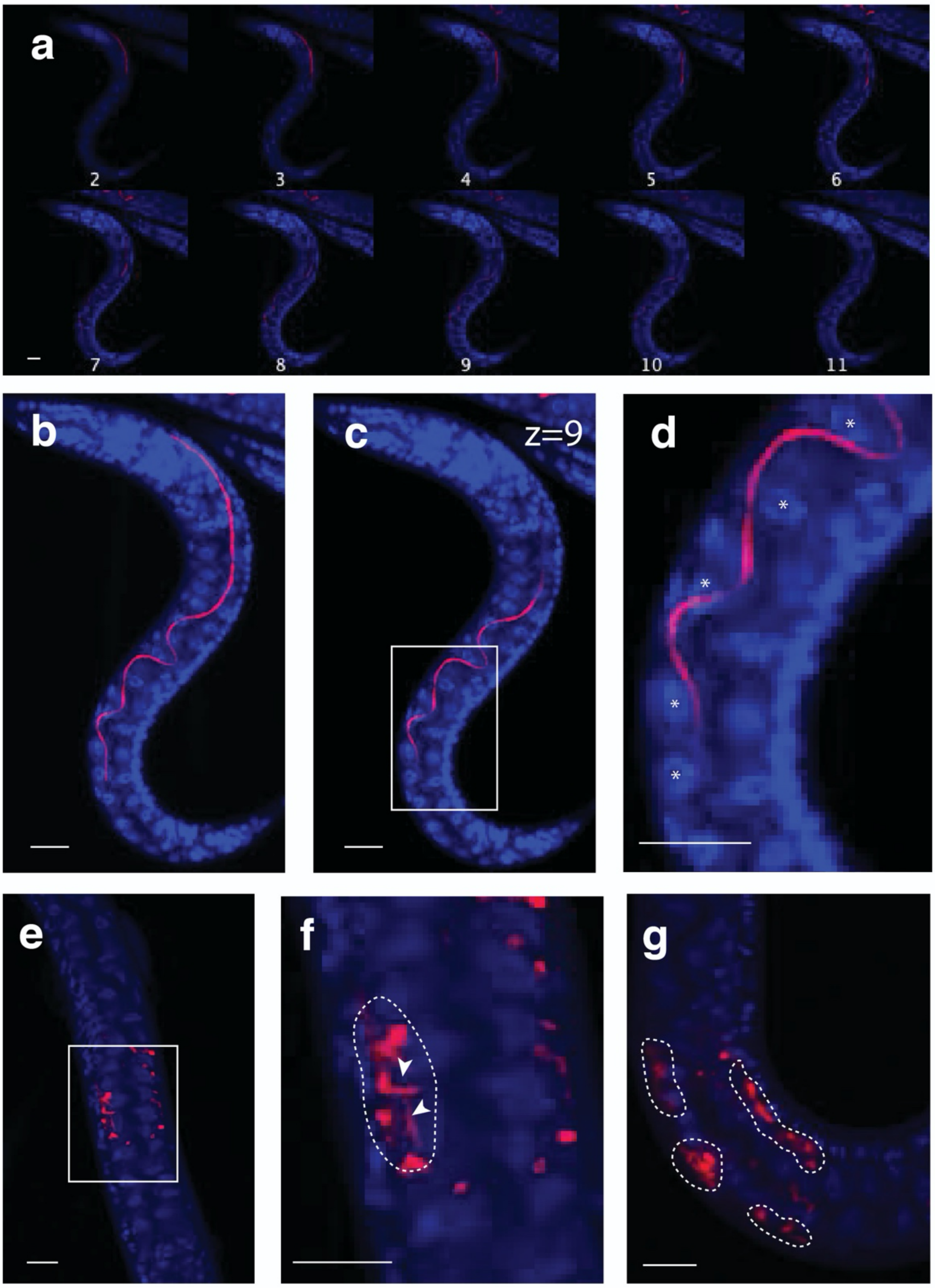


Supplementary Fig. 6. **Contiguous infection events. a** A representative montage of an WT *B. atropi* infected animal with a distinguishable “filament” infection event (numbers indicate z planes). **b** Z-projection image of the animal in **a** showing the full length of a filament. **c** Representative plane 9 showing a filament passing by multiple intestinal cells. **d** Zoomed-in region in the white box in **c** with counted intestinal nuclei indicated by asterisks. **e** An example of a LUAb7-infected animal with an infection focus. **f** Inset of region indicated in white box in **e** showing short filaments (arrowheads) closely spaced to one another and nearby coccobacilli assumed to originate from a single infection event. **g** An example of an animal with multiple infection foci (dashed line). Scale bars are 10 μm.

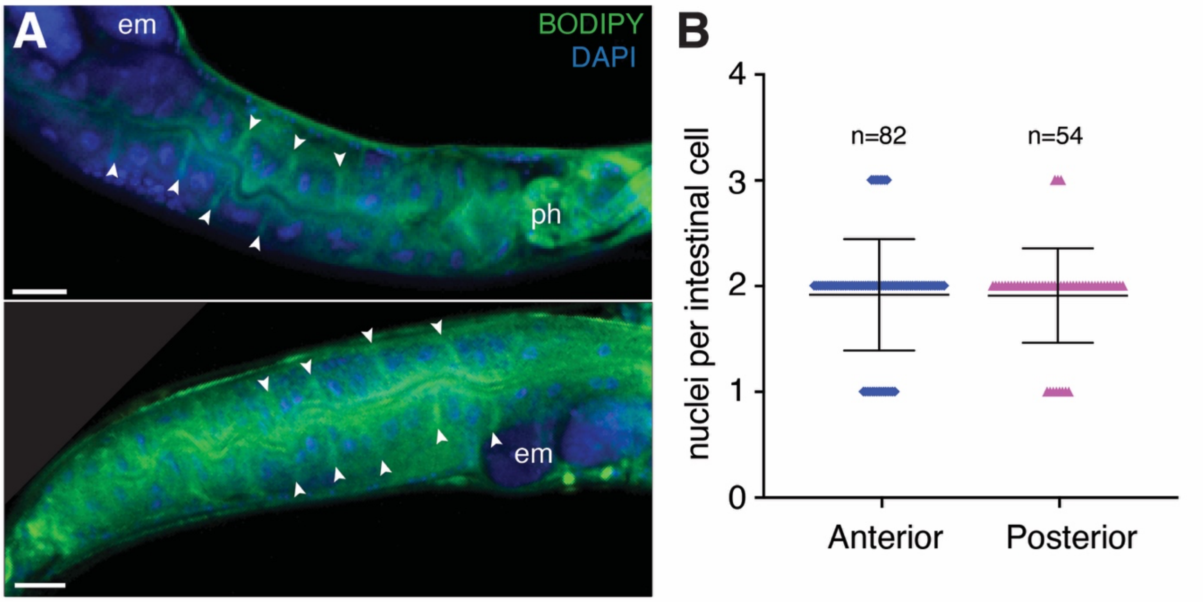


Supplementary Fig. 7. ***O. tipulae* intestinal cells contain an average of 2 nuclei at the anterior and posterior. a** Confocal images of *O. tipulae* animals stained with mixture of CellBrite, BODIPY-ceramide and DAPI, with lateral intestinal membranes indicated (*arrowheads*), as well as the pharynx (*ph*), and embryos (*em*). Scale bars are 20 μm. **b** The number of nuclei in intestinal cells with distinctly stained lateral membranes was counted at the anterior half and posterior half of several animals. The mean (1.9 for both) and SD are shown.

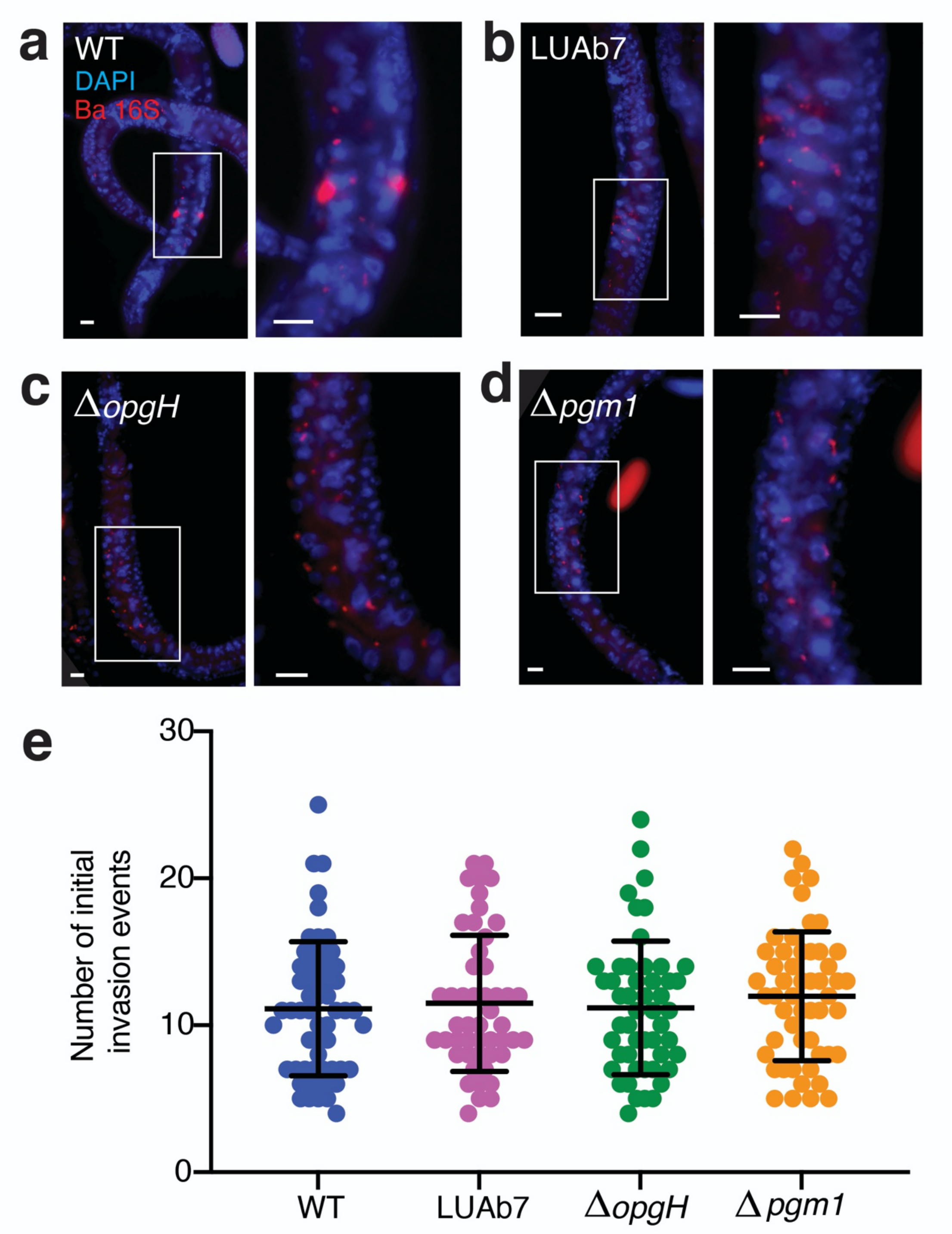


Supplementary Fig. 8. **Short pulse-chase infection results in similar initial invasion events across different strains at 16 hpi. a-d** Representative images of animals infected with WT, LUAb7, or knockout strains at 16 hpi showing similar numbers of invading bacteria. White boxes indicate regions of interest examined at higher magnification. Scale bars are 5 μm. **e** Quantification of **a-d**. Results are from 2 independent replicates, n > 47 animals per group.

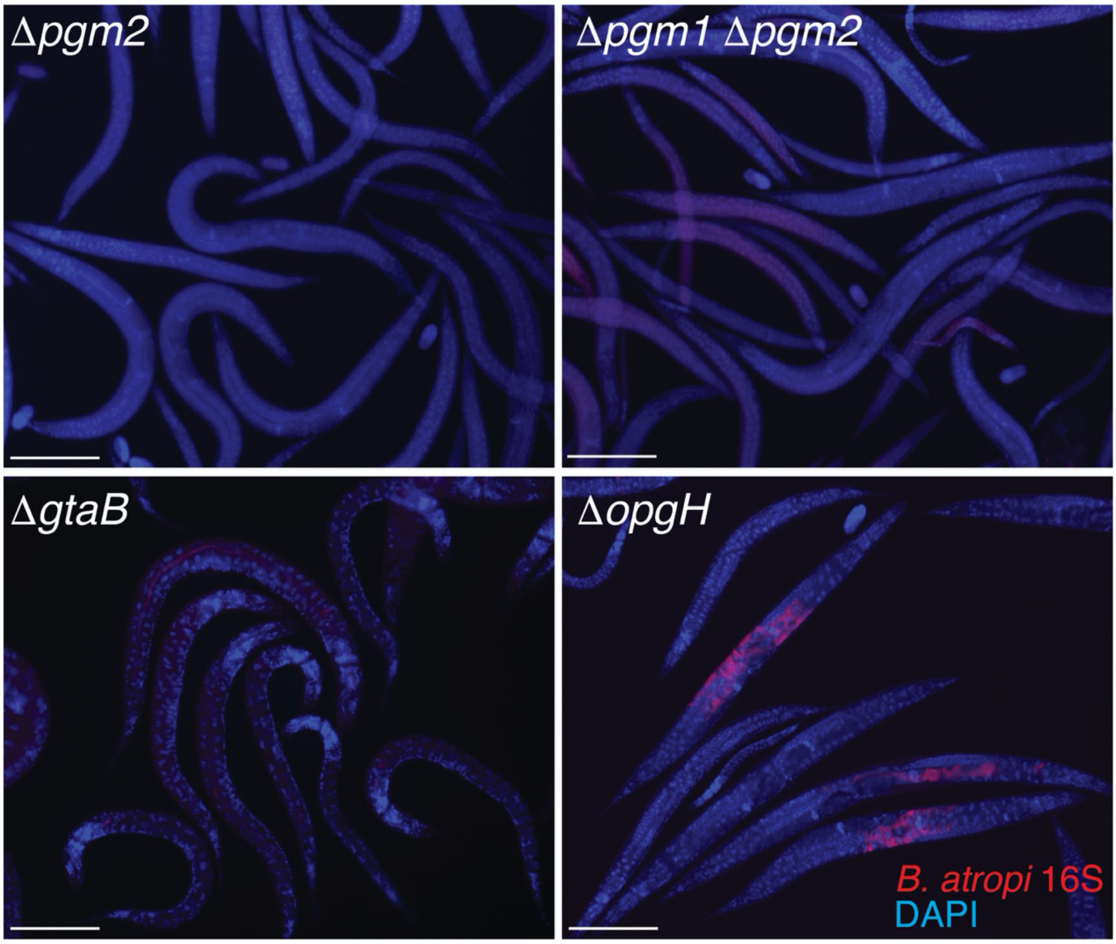

Supplementary Fig. 9. **Lack of in vivo infection in *pgm2* and *gtaB* knockout mutants.** JU1501 *O. tipulae* animals were pulse infected for 2 hours with indicated *B. atropi* strains and harvested 34 hpi for staining with 16S FISH and DAPI. Scale bars are 50 μm.
